# Burn-induced decreases in soil microbial carbon use efficiency vary across soil types and substrates

**DOI:** 10.1101/2025.07.31.667753

**Authors:** Dana Johnson, Kara Yedinak, Thea Whitman

## Abstract

Wildfires cause immediate changes in above and belowground carbon (C) stocks in boreal forest ecosystems with long-term repercussions for C cycling. Understanding the role of soil microbes in mediating post-fire C cycling and recovery is an important step to predicting how these ecosystems will respond to novel wildfire regimes caused by climate change. Wildfires can cause large shifts in soil bacterial and fungal community composition that can persist for years post-fire. Less is known about the effects of fire on soil microbial community function, such as C use efficiency (CUE). In this study, we measured the effects of burning on substrate-specific CUE using a laboratory incubation of boreal forest soils. We amended burned and unburned soils with either ^13^C-labelled ground pine roots or glucose and measured the amount of added substrate C that was incorporated into microbial biomass C versus respired as CO_2_ in order to calculate CUE. Burning caused a decrease in the amount of soil microbial biomass and respiration derived from soil organic C. Glucose-specific CUE declined with burning, driven by a decrease in glucose-derived microbial biomass. This decrease in glucose-specific CUE following burning correlated with an increase in weighted mean predicted 16S rRNA gene copy number, raising the possibility of using copy number as a proxy for post-fire CUE in boreal forest soils. Overall, pine-specific CUE was lower than glucose-specific CUE, likely reflecting the difference in chemical complexity between the two substrates; burning had a much smaller effect on pine-specific CUE, highlighting the variability of CUE between substrates in burned soils.

## 1. Introduction

Boreal forests hold a globally important reservoir of carbon (C) both above- and belowground (Bradshaw and Warkentin 2015). Understanding how climate change and wildfires will impact boreal forest C cycling requires knowledge of controls on C stocks and decomposition rates. A growing number of studies document shifts in soil microbial community composition following fire (*e.g.*, Holden et al. 2016, Pérez-Valera et al. 2019, Whitman et al. 2019, Dove et al. 2022, Johnson et al. 2023, 2024). However, compositional shifts may or may not correspond to shifts in function, and genome-based analyses only allow us to infer predicted functional potential (Cobo-Díaz et al. 2015, Dove et al. 2022, Nelson et al. 2022). Directly assessing the functions of soil microbes will allow us to better understand whether and how microbially mediated processes change soil organic C (SOC) cycling post-fire.

One critical way wildfires could affect microbial community function is via changes in carbon use efficiency (CUE), which may have important implications for post-fire soil C stocks as was previously posited by Nelson *et al*. (2024). CUE is an estimate of the proportion of total substrate C taken up by microbes that is converted into microbial biomass C. There are numerous methodologies for measuring microbial community CUE, involving ^13^C-labelled substrate tracing (Frey et al. 2013), ^18^O-water tracing (Blazewicz and Schwartz 2011), calorespirometry (Hansen et al. 2004, Wadsö 2009), metabolic flux analysis (Dijkstra et al. 2011), and even inference from genomic data (Saifuddin et al. 2019), which all have strengths and limitations (see Geyer et al. 2019 and He et al. 2024 for a comprehensive discussion).

CUE is affected by available C substrates, microbial community composition, and environmental conditions – all of which can be altered by fire (Figure 1). For example, because more complex C compounds, such as cellulose, generally require more enzymatic steps for degradation than less complex C substrates, such as glucose, CUE generally decreases with increasing complexity of the C substrate (Frey et al. 2013, Öquist et al. 2017). This may be of particular relevance in the post-fire soil environment due to the heat-induced conversion of organic matter to pyrogenic organic matter. Pyrogenic organic matter has a higher degree of aromaticity than unburned organic matter (Wiedemeier et al. 2015), thus, we might expect the CUE of pyrogenic organic matter to be lower than that of its unburned predecessor. Fire may also affect CUE via shifts in the C:N ratios of remaining C substrates. Because the temperature threshold for the complete volatilization of C is lower than that of N, fire can cause a decrease in C:N ratios in surface soils where burning temperatures are highest (Araya et al. 2017, Johnson et al. 2023, 2024). Negative correlations between CUE and C:N ratios have been observed in incubated soils (Manzoni et al. 2012), raising the possibility that fire-induced decreases in C:N ratios could increase CUE.

**Figure 1.**
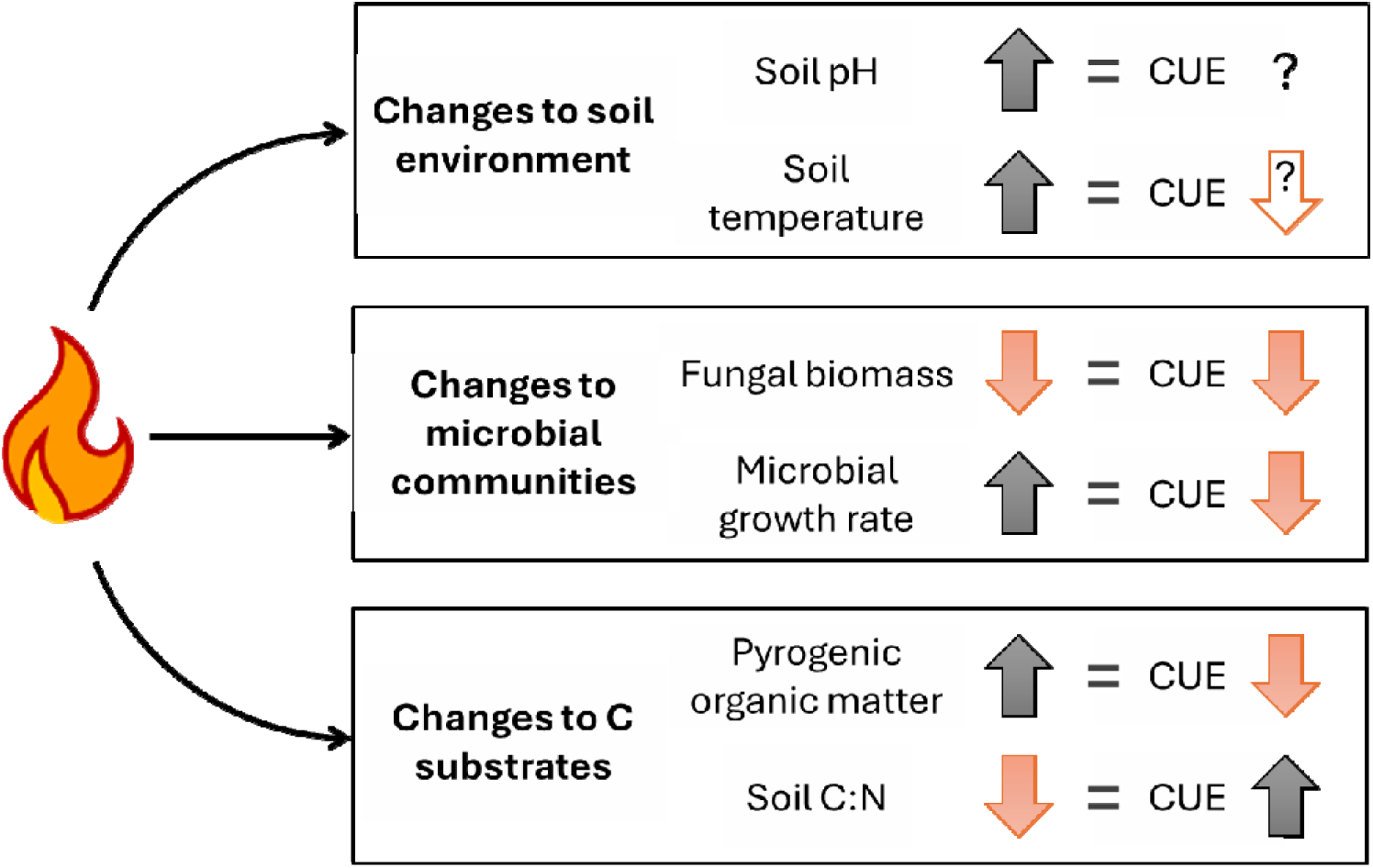
Hypothesized mechanisms by which fire could affect carbon use efficiency (CUE) via changes in the soil environment, microbial communities, and C substrates.

In addition to altered CUE post-fire due to changes in substrate, CUE could also be affected by shifts in microbial communities towards taxa with lower CUE as suggested by Nelson *et al*. (2024). For example, fire-induced changes in the ratio of fungal to bacterial biomass could affect CUE. Fires typically cause larger reductions in fungal biomass than bacterial biomass (Pressler et al. 2019), and fire has been observed to cause a decrease in fungi:bacteria ratios in boreal forest surface soils (Zhou et al. 2019). Fungi are thought to have higher CUE than bacteria, as has been measured in fungal and bacterial isolates (Lipson et al. 2009, Keiblinger et al. 2010). Thus, higher fire-induced fungal mortality could contribute to decreased CUE.

Another way in which shifts in microbial community composition may affect CUE is via changes in microbial growth rates. Fire causes heat-induced microbial and plant mortality, and the resulting necromass is a source of nutrients that may open a niche for fast-growing taxa in the soil. In boreal forest soils, fast-growing taxa were observed to be enriched in burned soil compared to unburned soil one year following wildfire (Johnson et al. 2023). An increase in the relative abundance of fast-growing taxa and the resulting increase in mean microbial community growth rate may drive a decrease in CUE due to a tradeoff between growth rate and yield, as has been previously observed (Pfeiffer et al. 2001, Lipson et al. 2009). In studies of bacterial isolates, Roller et al. (2016) found that CUE is negatively correlated with maximum potential growth rate and 16S rRNA gene copy number. Because is it difficult to directly measure growth rates of a soil microbial community, predicted weighted mean 16S rRNA gene copy number has been used as a proxy for growth rate (Yano et al. 2013, Roller et al. 2016), and increased mean 16S rRNA gene copy numbers in forest soils have been observed in the aftermath of fires (Nemergut et al. 2015, Whitman et al. 2019, Johnson et al. 2023). While it has been hypothesized that CUE should decrease with increasing rRNA gene copy number, there is very little data on this relationship. One previous study using bacterial isolates from a temperate deciduous forest found a positive relationship between rRNA gene copy number and CUE (Pold et al. 2020). In a second laboratory study focused on litter additions and manipulations of soil minerology, researchers documented a decrease in CUE with both increasing rRNA gene copy number and maximum microbial growth rate (Elias et al. 2024). In this study, we present the first data on the relationship between CUE and predicted weighted mean rRNA gene copy number following wildfire.

Microbial CUE may also be affected by environmental conditions such as soil pH or temperature. For example, burning can cause large increases in soil pH (Certini 2005, Úbeda et al. 2005, Johnson et al. 2023), which may influence CUE, but the relationship between CUE and soil pH is complicated. CUE has been observed to be lowest in soils at circumneutral pH and higher in soils with pH farther from circumneutral across a wide range of ecosystems (Sinsabaugh et al. 2016, Schroeder et al. 2024). However, in a study of more than 900 agricultural soils across Western Australia, researchers observed lower CUE values in acidic soils (pH 4) and highest CUE in soils with pH ranging from 5.5 to 7.5 (Jones et al. 2019). There is very little data on the relationship between CUE and soil pH specific to boreal forest ecosystems. In one study of forest soils from Sweden, increasing soil pH via liming was observed to cause an increase in CUE (Silva-Sánchez et al. 2019); thus, while it is possible that fire-induced increases in soil pH in acidic soils may contribute to lower CUE values, we expect that additional factors not yet fully understood may affect the relationship between pH and CUE in boreal forest soils.

Changes in soil temperature in the post-fire environment may also contribute to lower CUE. Some studies have shown a decrease in CUE with increasing temperature (Manzoni et al. 2012), although similar to pH, the relationship between CUE and soil temperature is not consistent across studies. For example, a recent metanalysis found no consistent relationship between soil warming and CUE (Zhang et al. 2024). Ultimately, the relationship between CUE and temperature is likely nuanced. For example, Frey et al. found that microbial CUE was dependent on both soil temperature and substrate quality, with the CUE of glucose unaffected by increasing temperature while the CUE of more complex substrates declined (Frey et al. 2013). Following wildfire, soil temperature can be elevated by several degrees due to the lack of vegetation cover, loss of the organic surface layer, and increased surface albedo driven by pyrogenic organic matter and ash deposition (Burke et al. 1997, O’Neill et al. 2002, Treseder et al. 2004, Kasischke and Johnstone 2005, Kelly et al. 2021). The net effect of increased soil temperature post-fire on CUE would be expected to be modulated by the availability of various C substrates. While the ultimate effect of burning on soil CUE is likely affected by a combination of all these factors – available C substrates, microbial community composition, and environmental conditions – the collective predictions for CUE after fire tend to be negative (Figure 1), which is consistent with observations of decreased CUE in burned forest soils (Auwal et al. 2023). The effect of fire on CUE is likely also shaped by fire characteristics such as burn severity and would be expected to change over time.

Disentangling the impact of fire across different severities on soil microbial CUE is particularly important for improving model-based predictions of post-fire C emissions. C models are an important tool in the work towards understanding the effects of ecosystem disturbances, such as wildfire, on soil C stocks, and CUE is a common constant used in C cycling models (Cotrufo et al. 2013, Sulman et al. 2018). There is strong interest in integrating shifts in microbial community functioning, such as maximum enzymatic rate (V_max_) and CUE, into existing C models to better study how disturbance events affect microbial functioning and soil C cycling. However, we were only able to find one study that measured the impact of wildfire on soil CUE (Auwal et al. 2023). Currently, CUE is typically represented as a constant in C models, precluding any fire-induced shifts in CUE or return of CUE to pre-burn levels. Increasing our understanding of CUE in post-fire soils will allow more accurate models of microbially mediated C cycling following wildfires.

Here we investigate how burning and varying burn duration impact microbial community function as measured by changes in substrate-specific CUE in boreal forest soils. We measured post-fire pine- and glucose-specific CUE in both organic-rich (Histosols) and sandy soils (Gleysols) characteristic of boreal forests in north-central Canada. We hypothesized that:

(1) Burning will cause CUE to decline in both Histosols and Gleysols. We predicted this because we expect shifts in microbial communities towards faster-growing taxa.

(2a) Pine-specific CUE (CUE_pine_) will be lower than glucose-specific CUE (CUE_glucose_) because pine wood is composed of more complex molecules than glucose.

(2b) Substrate-specific differences (pine *vs*. glucose) will be smaller than changes in CUE driven by burning. Our rationale was that the proliferation of fast-growing microbial taxa post-fire would overshadow the effects of substrate.

(2c) The difference in substrate-specific CUE in Gleysols will be smaller than in Histosols. We predicted this because the sandy soil cores were collected from sites dominated by pine trees, and thus, these microbial communities may have a “home field advantage” at degrading pine (Fanin et al. 2016).

## 2. Methods

### 2.1. Soil collection and burns

We used laboratory burns to simulate boreal crown fires on intact soil cores collected from soils within Wood Buffalo National Park, Alberta, Canada, to measure the effects of burning on soil microbial CUE. We randomly identified 12 sites across the park that represented two dominant vegetation types – 6 jack pine sites (*Pinus banksiana* Lamb.) and 6 spruce sites (*Picea mariana* (Mill.) and/or *Picea glauca* (Moench) Voss) – and were from areas of the park that had not burned for more than 30 years (Table S1). The *Pinus banksiana*-dominated sites were underlain by soils classified as Eutric Gleysols, and the *Picea* spp. sites were underlain by soils classified as Dystric Histosols (Food and Agriculture Organization of the United Nations, 2003). At each site, for a connected series of studies (Johnson et al., 2024), we collected 16 soil cores 6 cm in diameter and 10 cm in height (Table S2), three of which were randomly selected for the CUE experiments described in this paper. The soil cores were transported to Madison, WI, USA within 10 days of sampling and then air dried for 6 weeks to simulate drought conditions.

Laboratory burns were conducted by exposing pairs of intact soil cores from each sampled site to a 60 kW m^-2^ heat flux in a Mass Loss Calorimeter (Fire Testing Technology Limited, West Sussex, UK) for either 120 s or 30 s (see Johnson et al. 2024 for further details). During and following the burns, beaded 20-gauge Type-K thermocouples (GG-K-20-SLE, Omega) were used to track soil temperatures at the midpoint of the core (either at the base of the O horizon in Gleysol cores or 5 cm below the soil surface in Histosols) and 1 cm above the base of the core. A third core from each sampled site was not burned (designated “control” cores). Following the laboratory burn, water was added to each intact core to bring moisture to 65% of field capacity and cores were placed in 475 mL Mason jars to incubate in the dark at room temperature. Jars were capped to limit moisture loss and opened for 5 minutes daily to prevent the development of anoxic conditions. Cores were incubated for 24 days before measuring CUE. This duration was chosen to allow soil microbial communities to begin to recover from the effects of burning and is more representative of intermediate-term effects than assessing microbial community dynamics immediately after the disturbance.

### 2.2. ^13^C amendments and tracing

After the 24-day incubation, the intact soil cores were destructively sampled. O and mineral horizons were separated and subsamples were collected for measurement of substrate-specific CUE using ^13^C-labelled glucose and pine following the ^13^C-tracing method from Geyer et al. (2019). Four subsamples were collected from each soil horizon and placed in 125 mL Mason jars. For Histosols, 1.5 g of soil was used, for Gleysol organic horizons, up to 2.0 g of soil was used, and for Gleysol mineral soils, 10 g of soil was used. The mass of soil used was limited by total intact horizon mass. Thus, for six of the 18 soil horizons, less than 2.0 g of organic soil (0.7-1.3 g) was used. ^13^C-labelled glucose (5 at%) or ^13^C-labelled pine (1.6 at%) was then added to two of the jars and unlabeled glucose (1.1 at%) or pine (1.1 at%) was added to the remaining two jars at a rate of 1.0 g glucose g^-1^ dry soil (0.4 mg C g^-1^ dry soil) or 2.4 mg pine g^-1^ dry soil (0.91 mg C g^-1^ dry soil), respectively (Table S3), and soil was mixed well. These rates were chosen with the goal of achieving detectable respiration and incorporation into microbial biomass while minimizing unrealistically large perturbances to the community due to new substrate additions. The pine amendment was created by finely grinding pulse-labelled roots of eastern white pine (*Pinus strobus* (L)). Briefly, we used the belowground biomass from two-year-old *Pinus strobus* tree seedlings grown in two custom growth chambers for one growing season under controlled moisture, humidity, temperature, and light conditions. In the first chamber, trees were pulse labeled with 99% ^13^CO_2_ at regular intervals (“labelled pine”). In the second chamber, trees were exposed to ambient levels of non-enriched CO_2_ (“unlabeled pine”) (full details in Zeba et al. 2024).

Jars were capped with lids fitted with rubber septa, sealed tightly, and incubated in the dark at 22 □. A gas syringe was used to collect 12 mL headspace gas samples every 24 hours, and gas samples were injected into 12 mL exetainers and diluted 1:1 with CO_2_-free air for later analysis. Gas samples were also collected from additional empty jars to measure background CO_2_ levels. Following sampling, jars were opened, allowed to equilibrate with external atmosphere for five minutes, re-capped, and incubated for another 24 hours. A total of four headspace gas samples were collected from each jar over a 4-day period. Production and isotopic signature of CO_2_-C of headspace gas samples was measured by analyzing C concentration and isotopic composition on a cavity ring-down spectrometer (SAM 22001 autosampler; G2201-isotopic analyzer for CO_2_ and CH_4_, Picarro).

The degree of ^13^C-labelled substrate incorporated into microbial biomass was measured using a simultaneous chloroform fumigation extraction modified from Gregorich et al. (1990). Briefly, soil subsamples were placed in 20 mL of 0.05 M K_2_SO_4_ solution to which 0.5 mL ethanol-free chloroform was added (“fumigated”). 10× less concentrated K_2_SO_4_ was used than the original method so that C concentration in the salt solution was sufficiently high so as to be detectable upon combustion (Potthoff et al. 2003, Makarov et al. 2015, Pang et al. 2021). Duplicate samples with no chloroform were also used (“non-fumigated”). Solutions were shaken at 150 revolutions per minute for four hours after which solutions were filtered at 45 µm using a syringe filter and resulting dissolved organic carbon (DOC) was collected and stored. The DOC-K_2_SO_4_ solution was dried down at 80 □ for 24 hours, and the C concentration and isotopic composition of the resulting salts were quantified on a Thermo Delta V Isotope ratio mass spectrometer (IRMS) interfaced to a NC2500 elemental analyzer at the Cornell University Stable Isotope Laboratory.

### 2.3. Calculation of CUE

Total microbial biomass C (MBC; µg C g^-1^ dry soil) was calculated using Equation 1 (see supplementary information for full list of equations and derivations):

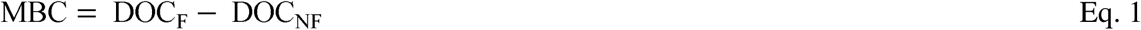

where DOC is the total C concentration of fumigated (F) and non-fumigated (NF) K_2_SO_4_ extracts on a per g dry soil basis. We did not correct MBC using an extraction efficiency coefficient (often termed k_EC_) because true extraction efficiencies vary widely across different soil properties (Dictor et al. 1998) and communities (Tate et al. 1988). Rather than multiplying all MBC estimates by the same value, which is almost certainly incorrect, we acknowledge that true MBC (and, hence, derived CUE) is presumably higher than our estimates by an unknown factor. Because we are primarily interested in comparing CUE from samples collected from the same site, which should have similar extraction efficiencies, this decision should not affect the direction of any observed trends.

Total microbial incorporation of ^13^C-labelled substrate C (MBC_LA_; µg C g^-1^ dry soil) was calculated as the product of MBC and the fraction of MBC that is substrate-derived (*f*_MBC,LA_) using Equations 2-5:

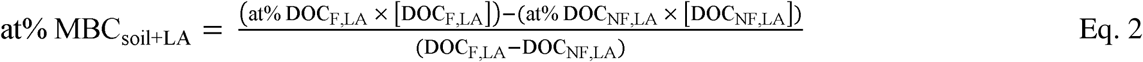

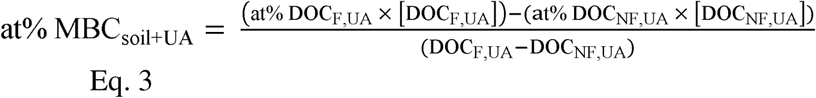

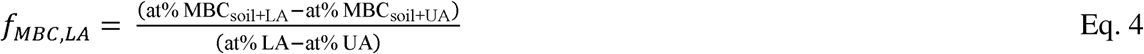

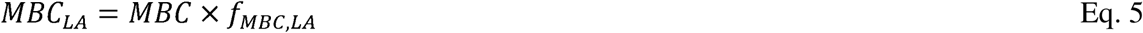

where at% MBC_soil_ and at% DOC represent the atom % of ^13^C in MBC and K_2_SO_4_ extracts, and DOC represents the concentration of C (µg C g^-1^ dry soil) of K_2_SO_4_ extracts from soils amended with labelled (LA) and unlabeled (UA) substrate, that were fumigated (F) or non-fumigated (NF). At% LA and at% UA are the atom % of ^13^C in the labelled and unlabeled substrates, respectively.

The total CO_2_ derived from the ^13^C-labelled substrate (CO_2,LA_; µg C g^-1^ dry soil) was calculated as the product of cumulative CO_2_-C respired (CO_2,total_; µg C g^-1^ dry soil) and the fraction of total CO_2_-C that is derived from the ^13^C-labelled substrate (*f*_CO2,LA_) using Equations 6-7:

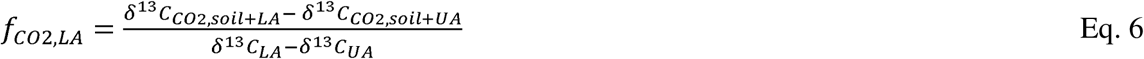

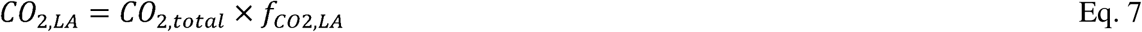

where δ^13^C_CO2,soil+LA_ and δ^13^C_CO2,soil+UA_ represent the δ^13^C values of CO_2_ from soils amended with the labelled (LA) or unlabeled (UA) substrate, respectively, and δ^13^C_LA_ and δ^13^C_UA_ represent the δ^13^C values of the labelled and unlabeled substrates, respectively.

Due to our relatively subtle addition of labelled substrates, substrate-derived MBC was not always detectable. Thus, before calculating CUE, we first filtered the data to include only datapoints with a positive value of substrate-derived MBC, which reduced the number of samples for pine-amended soils from 54 to 45 and for glucose-amended soils from 54 to 53. Of the 10 datapoints removed, 9 were for burned soil. CUE was then calculated using equation 8:

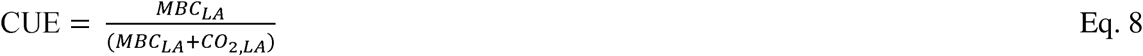

### 2.4. Estimating weighted mean 16S rRNA gene copy number

To assess the relationship between CUE and 16S rRNA gene copy number, we extracted DNA from soil 24 days post-burn using DNeasy PowerLyzer PowerSoil kits (QIAGEN) following manufacturer’s instructions. The 16S rRNA gene v4 region of bacteria and fungi was amplified via triplicate PCR using 515f and 806r primers (Walters et al. 2015) with added barcodes and Illumina sequencing adapters (Kozich et al. 2013). PCR amplicon triplicates were pooled, purified, and normalized using a SequalPrep Normalization Plate Kit (ThermoFisher Scientific, Waltham, MA), and then all samples were combined. Library cleanup was performed using a Wizard SV Gel and PCR Clean-Up System (Promega, Madison, WI). The pooled library was sequenced using 2 x 300 paired-end Illumina MiSeq sequencing at the UW-Madison Biotechnology Center, resulting in a total of 13M reads.

We quality filtered, trimmed, dereplicated, learned errors, picked OTUs, and removed chimeras using DADA2 (Callahan et al. 2016) implemented in QIIME2 (Bolyen et al. 2019), which resulted in the retention of 6.5M reads (mean 28,879 reads per sample). We excluded 3 samples from further analysis due to low 16S reads per sample (<1000). Taxonomy was assigned using the naïve Bayes classifier (Bokulich et al. 2018) in QIIME2 with the aligned 515f-806r region of the 99% OTUs from the SILVA database (SILVA 138 SSU) (Quast et al. 2013, Yilmaz et al. 2014, Glöckner et al. 2017).

To estimate mean predicted 16S rRNA gene copy number, we used the rrnDB RDP Classifier tool (v.2.14) to assigned a predicted mean 16S rRNA gene copy number for each genus in this study (Stoddard et al. 2015). For taxa without a genus-level assignment, we used the mean copy number for all taxa with a genus-level assignment in the study. We then calculated community-weighted mean predicted rRNA gene copy number by dividing OTU counts by the predicted copy number, calculating relative abundance of each OTU, and summing the product of the predicted copy number and each OTU’s relative abundance for all OTUs in each sample (Nemergut et al. 2015).

(We draw on this sequencing data in this paper only to explore its relationship to CUE at a single timepoint. An associated paper following bacterial and fungal community composition across four timepoints post-burn is in preparation.)

### 2.5. Statistical Analyses

We performed all analyses using R statistical software version 4.4.0 (R Core Team 2024) and plotted model results using R package *ggplot2* (Wickham 2016). We used ANOVA models including interaction terms between C amendment and burn treatment exposure and Tukey’s HSD to assess the impact of burning, burn treatment exposure, and soil type on MBC, CO_2_ respired, and substrate-specific CUE (Oksanen et al. 2020). We performed nonlinear (exponential) regressions to evaluate the relationships between substrate-specific CUE and maximum soil temperature, predicted weighted mean 16S rRNA gene copy number, and soil pH.

## 3. Results

### 3.1. Burning decreased SOC-derived microbial biomass and cumulative CO_2_ respired in organic horizon soils

Burning decreased both SOC-derived MBC and cumulative SOC-derived CO_2_ from organic horizon soil (Figure 2). The small masses of added substrates (Table S3) had little overall effect, as desired: the bulk of MBC and CO_2_ emitted over the four day incubation were derived from SOC and not from the added glucose or pine, and substrate additions did not have a significant effect on SOC-derived CO_2_ respired (p = 0.97), nor was the interaction between substrate (*i*.*e*., glucose *vs*. pine) and burn treatment significant (p = 0.99).

**Figure 2.**
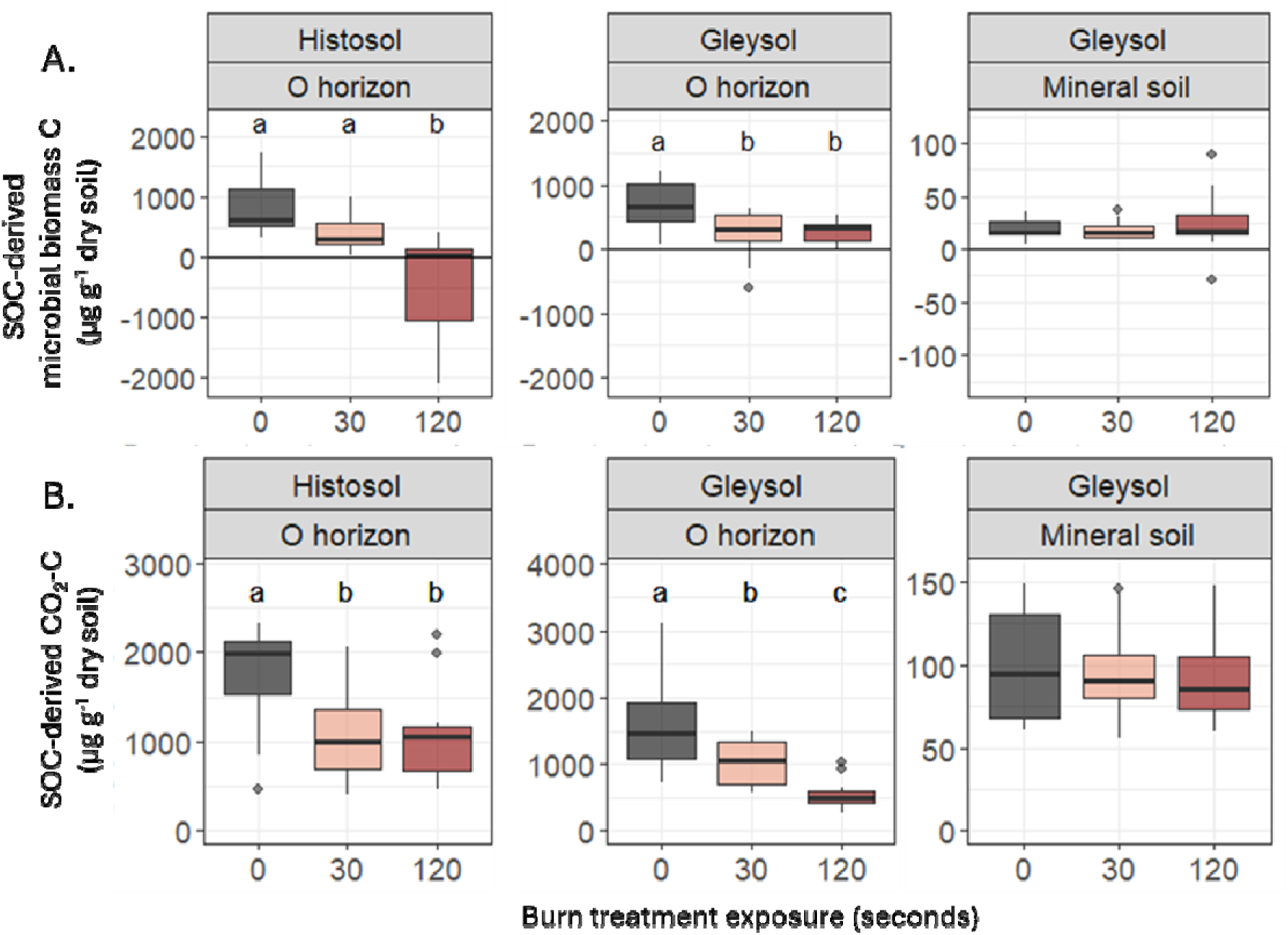
SOC-derived microbial biomass C (A) and cumulative SOC-derived CO_2_-C over the four-day incubation (B) in Histosols (left panels), Gleysol organic horizons (middle panels) and Gleysol mineral soil (right panels) (N=108 total; n=6 for each boxplot). Different letters represent statistically significant differences in treatments based on ANOVA and Tukey’s HSD (p < 0.05). The central horizontal line indicates the median, the upper and lower bounds of the box indicate the inter-quartile range (IQR), the upper and lower whiskers reach the largest or smallest values within a maximum of 1.5 * IQR, and data beyond the whiskers are indicated as individual points. Substrate did not have a significant effect on microbial biomass or cumulative CO_2_-C respired, so glucose and pine-amended treatments are plotted together. Note different scales on y-axes.

SOC-derived MBC was higher in unburned Histosols (mean = 857 ± 503 μg C g^-1^ dry soil) than in Histosols exposed to the 120 s heat dose treatments, where it was statistically undetectable (mean = -430 ± 902 μg C g^-1^ dry soil, p < 0.001), but there was no significant difference following the 30 s heat dose treatments (mean = 404 ± 296 μg C g^-1^ dry soil, p = 0.2; Figure 2A). SOC-derived MBC was higher in the organic horizons of unburned Gleysols (mean = 674 ± 386 μg C g^-1^ dry soil) than in the organic horizons of Gleysols exposed to either the 30 s (mean = 247 ± 395 μg C g^-1^ dry soil, p = 0.002) or the 120 s heat dose treatment (mean = 282 ± 185 μg C g^-1^ dry soil, p = 0.04).

There was no significant difference in SOC-derived MBC in the mineral soil of unburned Gleysols (mean = 19 ± 9.7 μg C g^-1^ dry soil) *vs*. burned mineral soil (30 s exposure, mean = 18 ± 8.7 μg C g^-1^ dry soil, p = 1.0; 120 s exposure, mean = 24 ± 29 μg C g^-1^ dry soil, p= 0.8). Substrate did not have a significant effect on MBC (p=0.4), nor was the interaction between substrate (*i*.*e*., glucose *vs*. pine) and burn treatment significant (p=0.4).

Burning caused a decrease in SOC-derived CO_2_ respired from organic horizon soils, but there was no significant effect of burning on cumulative SOC-derived CO_2_ from mineral soils (Figure 2B). SOC-derived CO_2_ was higher from unburned Histosols (mean = 1723 ± 581 μg C g^-1^ dry soil) than in Histosols exposed to the 30 s (mean = 1082 ± 528, p = 0.003) or 120 s heat dose treatments (mean = 1077 ± 537 μg C g^-1^ dry soil, p = 0.003). SOC-derived CO_2_ was higher in the organic horizons of unburned Gleysols (mean = 1595 ± 712 μg C g^-1^ dry soil) than in mineral soil (unburned, Gleysol, mean = 98 ± 33 μg C g^-1^ dry soil) or the organic horizons of Gleysols exposed to either the 30 s (mean = 1012 ± 347 μg C g^-1^ dry soil, p = 0.005) or the 120 s heat dose treatment (mean = 555 ± 227 μg C g^-1^ dry soil, p < 0.001).

### 3.2. Microbial communities in burned soils incorporated less glucose-C into biomass but pine-C incorporation did not differ

Burning decreased the amount of amended glucose C incorporated into microbial biomass in organic horizon soils (Figure 3B). Glucose-derived MBC was higher in unburned organic horizon soils (Histosol, mean = 46 ± 15 μg C g^-1^ dry soil; Gleysol, mean = 50 ± 14 μg C g^-1^ dry soil) than in organic horizon soils exposed to the 30 s (Histosol, mean = 28 ± 7.7 μg C g^-1^ dry soil, p = 0.05; Gleysol, mean = 16 ± 11 μg C g^-1^ dry soil, p = 0.001) or 120 s heat dose treatments (Histosol, mean = 10 ± 10 μg C g^-1^ dry soil, p = 0.001; Gleysol, mean = 13 ± 5.9 μg C g^-1^ dry soil, p = 0.001).

**Figure 3.**
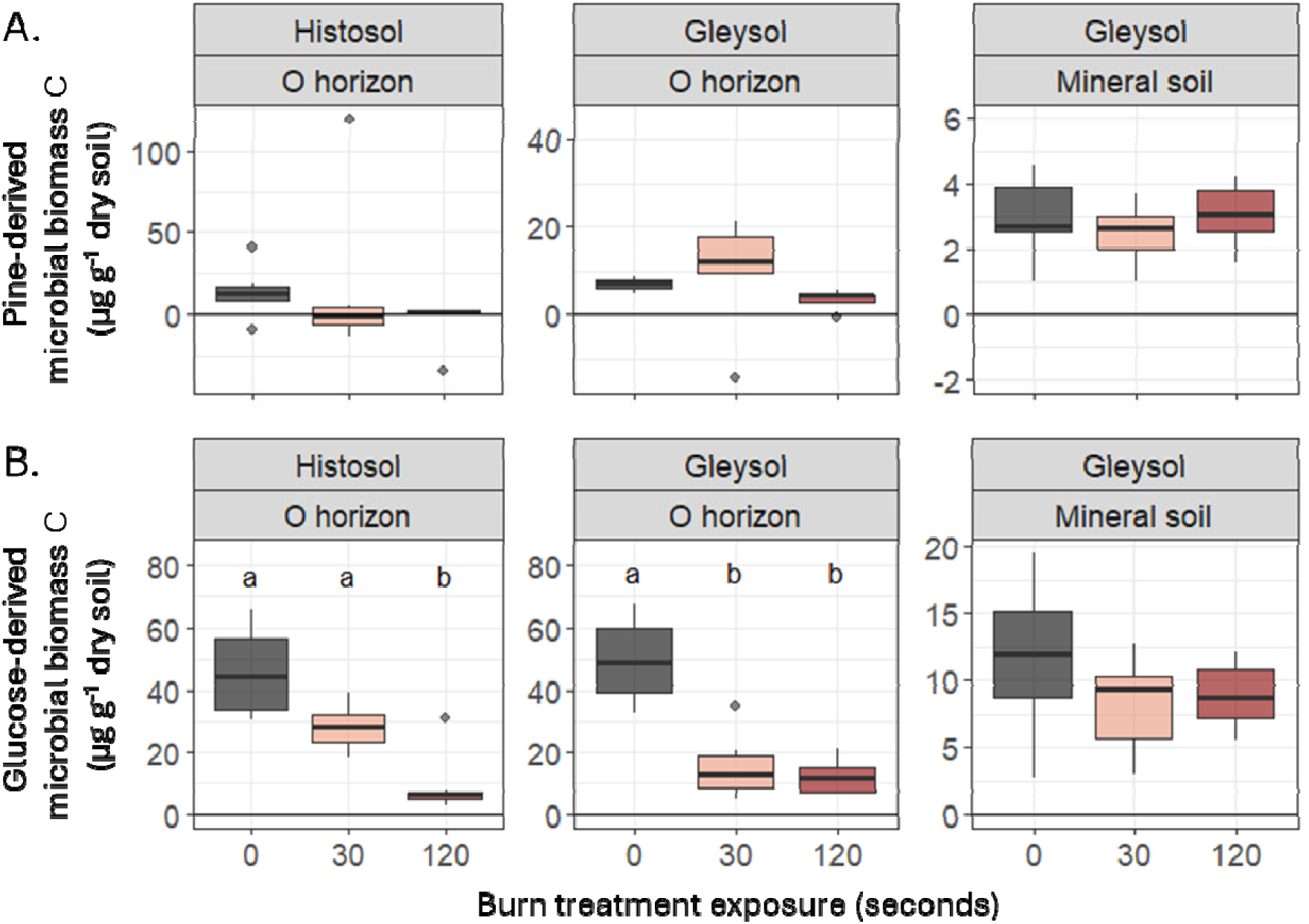
Microbial biomass C derived from (A) pine amendment and (B) glucose amendment in Histosols (left panel), Gleysol organic horizons (middle panel) and Gleysol mineral soil (right panel) (N=108 total; n=6 for each boxplot). Different letters represent statistically significant differences in treatments based on ANOVA and Tukey’s HSD (p < 0.05). The central horizontal line indicates the median, the upper and lower bounds of the box indicate the inter-quartile range (IQR), the upper and lower whiskers reach the largest or smallest values within a maximum of 1.5 * IQR, and data beyond the whiskers are indicated as individual points. Note different scales on y-axes.

Overall, across both soil types, despite more pine C being added per gram of soil, less of the pine amendment C was incorporated into microbial biomass (unburned, Histosol, mean = 14 ± 16 μg C g^-1^ dry soil; unburned, Gleysol organic horizons, mean = 7.0 ± 1.5 μg C g^-1^ dry soil, representing 0.7-1.4% of added pine C) than the glucose amendment (unburned, Histosol, mean = 46 ± 15 μg C g^-1^ dry soil, p = 0.005; unburned, Gleysol organic horizons, mean = 50 ± 14 μg C g^-1^ dry soil, p < 0.001, representing 12-13% of added glucose C; Figure 3).

The effect of burning on the incorporation of C from pine amendments into MBC was not significant (Figure 3A). There was a slight but not significant decrease in mean pine-derived MBC between unburned Histosols (mean = 14 ± 16 μg C g^-1^ dry soil) and Histosol samples exposed to the 120 s heat dose treatment (mean = -4.6 ± 15 μg C g^-1^ dry soil; p=0.6). Similarly, in the Gleysols, there was a slight decrease in mean pine-derived MBC between the unburned organic horizons (mean = 7.0 ± 1.5 μg C g^-1^ dry soil) and organic horizons exposed to the 120 s heat dose treatment (mean = 3.3 ± 2.4 μg C g^-1^ dry soil; p = 0.8).

### 3.3. Burning did not cause a decrease in pine-derived or glucose-derived CO_2_

Although burning decreased total respiration in organic horizon soils (Figure 2B), there was no significant effect of burning on pine-derived CO_2_ from either Histosols or Gleysols (Figure 4A). There was a small increase in glucose-derived CO_2_ from the organic horizons of Gleysols exposed to the 120s heat dose treatment (mean = 135 ± 24 μg C g^-1^ dry soil) compared to the unburned Gleysol organic horizons (mean = 106 ± 19 μg C g^-1^ dry soil, p = 0.05; Figure 4B). There was no significant effect of burning on glucose-derived CO_2_ from Histosols.

**Figure 4.**
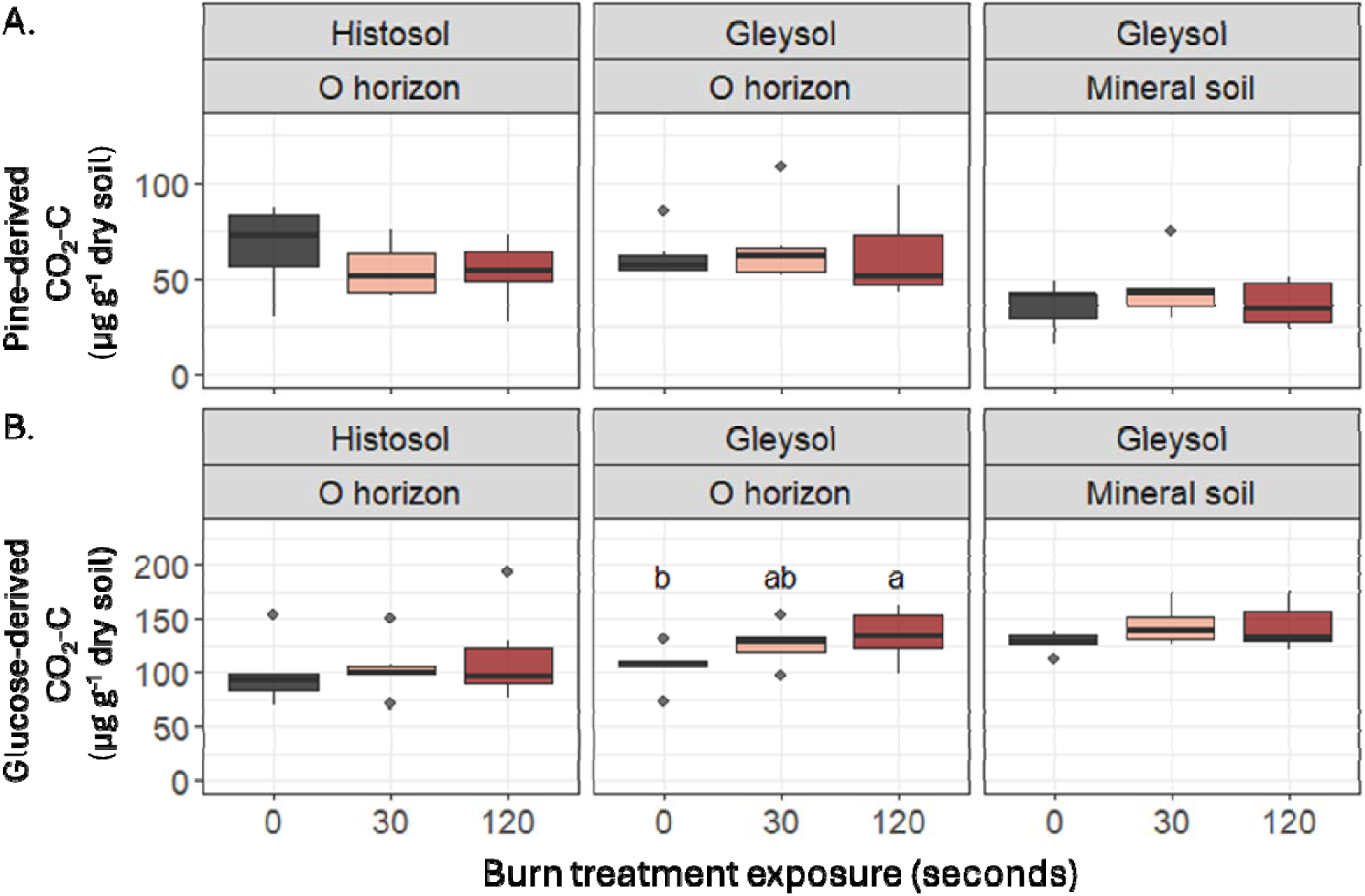
Cumulative CO_2_-C derived from either pine amendment (A) or glucose amendment (B) in Histosols (left panels), Gleysol organic horizons (middle panels) and Gleysol mineral soil (right panels) (N=108 total; n=6 for each boxplot). The central horizontal line indicates the median, the upper and lower bounds of the box indicate the inter-quartile range (IQR), the upper and lower whiskers reach the largest or smallest values within a maximum of 1.5 * IQR, and data beyond the whiskers are indicated as individual points.

Overall, substrate-derived CO_2_ was higher for glucose (mean = 122 ± 28 μg C g^-1^ dry soil, representing 31% of added glucose C) than pine (mean = 54 ± 19 μg C g^-1^ dry soil, p < 0.001, representing 5% of added glucose C).

### 3.4. Burning causes a large decrease in glucose-specific CUE and smaller effects on pine-specific CUE

CUE_glucose_ was not significantly different between Histosols and the organic horizon of Gleysols, and burning decreased CUE_glucose_ in organic horizon soils in both, with CUE_glucose_ values ranging from 0.02 in burned soil to 0.5 in unburned soil (Figure 5B). CUE_glucose_ was higher in unburned organic horizon (Histosol, mean = 0.3 ± 0.09; Gleysol, mean = 0.3 ± 0.1) than in organic horizon soils exposed to the 120 s heat dose treatments for both soil types (Histosol, mean = 0.09 ± 0.09, p = 0.001; Gleysol, mean = 0.08 ± 0.04, p = 0.001). For the 30 s heat dose treatment, CUE_glucose_ was significantly lower than in unburned organic horizons for Gleysols (mean = 0.1 ± 0.06, p = 0.001), but not significantly so for Histosols (mean = 0.2 ± 0.08, p = 0.1).

**Figure 5.**
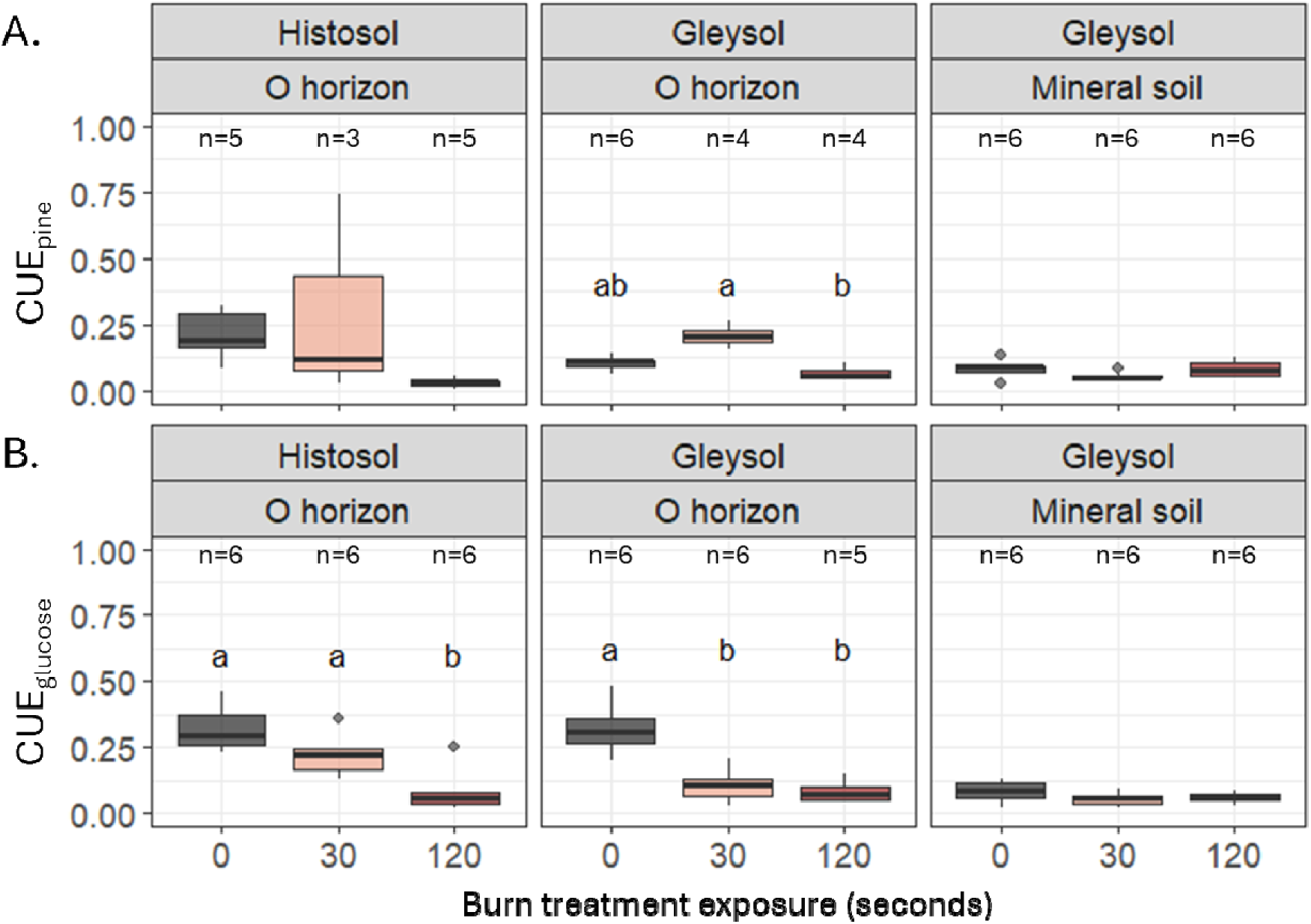
Microbial community (A) CUE_pine_ and (B) CUE_glucose_ in Histosols (left panels), Gleysol organic horizons (middle panels) and Gleysol mineral soil (right panels). Different letters represent statistically significant differences in treatments based on ANOVA and Tukey’s HSD (p < 0.05). The central horizontal line indicates the median, the upper and lower bounds of the box indicate the inter-quartile range (IQR), the upper and lower whiskers reach the largest or smallest values within a maximum of 1.5 * IQR, and data beyond the whiskers are indicated as individual points. Number of samples (n) is indicated for each boxplot.

CUE_pine_ was also not significantly different between Histosols and the organic horizon of Gleysols, with values ranging from 0.007 in burned soils to 0.3 in unburned soils (with one outlier of 0.7 in a burned Histosol; Figure 5A). However, the effects of burning on CUE_pine_ were generally not statistically significant: in the Histosols, exposure to the 120 s heat dose treatment decreased CUE_pine_ (mean = 0.03 ± 0.02) relative to unburned soils (mean = 0.2 ± 0.1), but the effect was not significant (p = 0.2). In the Gleysols, the only significant difference was in the organic horizons exposed to the 30 s heat dose treatment, which had significantly higher CUE_pine_ values (mean = 0.2 ± 0.04) than the organic horizons exposed to the 120 s heat dose treatment (mean = 0.07 ± 0.03, p = 0.04) and higher, albeit not significantly, CUE_pine_ values than the unburned organic horizons (mean = 0.1 ± 0.03, p = 0.06).

In the mineral soil of the Gleysol, there was no significant effect of substrate or burning on CUE, with CUE values ranging from 0.02 to 0.13.

There were small differences in CUE_pine_ *vs.* CUE_glucose_ across soil types and horizons. CUE_pine_ in unburned Histosols (mean = 0.2 ± 0.1) was slightly lower than CUE_glucose_ (mean = 0.3 ± 0.09, p = 0.08). In the organic horizon of Gleysols, CUE_pine_ (mean = 0.1 ± 0.03) was lower than CUE_glucose_ (mean = 0.3 ± 0.1, p < 0.001). There was no significant difference in CUE_pine_ *vs.* CUE_glucose_ in mineral soil.

We observed negative correlations between CUE_glucose_ and maximum soil temperature during the burn and soil pH. CUE_glucose_ decreased exponentially with increasing maximum soil temperature during burning in organic horizons (y = 0.6e^-0.04x^ + 0.07; p < 0.001, R^2^_adj._ = 0.62) and in mineral soil (y = 0.3e^-0.08x^ + 0.04; p = 0.06, R^2^ _adj_= 0.12; Figure 6A). CUE_glucose_ also decreased exponentially with increasing soil pH in Histosols but not Gleysols (y = 3.63^-0.41x^; p = 0.02, R^2^_adj._= 0.25 ; Figure S1). There was no significant correlation between CUE_pine_ and maximum soil temperature or soil pH.

**Figure 6.**
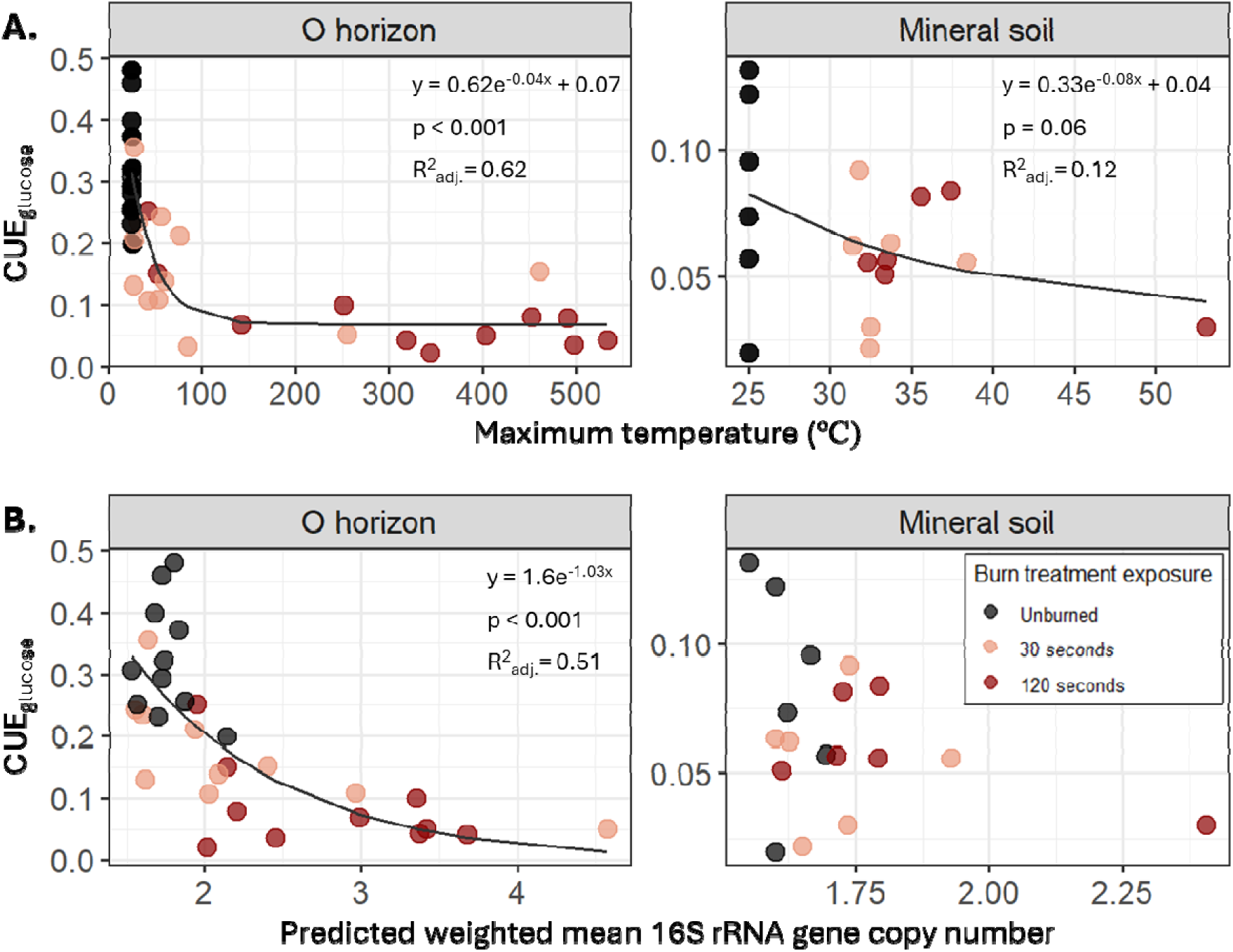
(A) Maximum soil temperature and (B) predicted weighted mean 16S rRNA gene copy number vs. CUE_glucose_ in organic horizon soils (left panels) and mineral soils (right panels). We used a non-linear regression to test for a significant correlation between weighted mean gene copy number and CUE. Soil type did not have a significant effect on the relationship between weighted mean gene copy number and CUE or between maximum temperature and CUE, so results for Histosols and Gleysols are plotted together.

### 3.5. CUE_glucose_ decreased exponentially with increasing 16S rRNA gene copy number

In organic horizons, there was an exponential decrease in CUE_glucose_ with increasing predicted weighted mean 16S rRNA gene copy number (Figure 6B; y = 1.6e^-1.03x^, p < 0.001, R^2^_adj_=0.51). There was not a significant relationship between CUE_pine_ and gene copy number (Figure S2). In the mineral horizons, there was no significant correlation between gene copy number and CUE_glucose_ or CUE_pine_.

## 4. Discussion

### 4.1. Burning decreases microbial biomass and respiration

Burning decreased both MBC and cumulative CO_2_ respired from organic horizon soils over the course of the four-day incubation (Figure 2) with little to no effect on underlying mineral soils. This is consistent with previous work documenting reductions in both soil microbial biomass and respiration following fire (Dooley and Treseder 2012, Pressler et al. 2019, Johnson et al. 2023, 2024). How long after a fire soil microbial biomass and respiration remain below pre-fire levels in boreal forest ecosystems will depend on numerous factors that govern these complex processes, such as the recovery of vegetation and the concomitant recovery of plant inputs to soil, which may take years to decades (Bond-Lamberty et al. 2004, Cuevas-González et al. 2009). In this study, our measurements of biomass and respiration were taken three weeks after the heat dose treatments, showing that, unsurprisingly, the effects of burning on microbial communities last at least weeks after fire.

While adding glucose or pine to our soils could have caused priming of existing SOC (Blagodatskaya and Kuzyakov 2008, Jilling et al. 2021), we observed no significant difference in SOC-derived microbial biomass or respired CO_2_ between soils amended with glucose *vs.* pine.

Although we did not have an unamended soil treatment to conclusively calculate any priming effects, given the difference in the complexity of these two C sources, we would not expect both substrates to induce priming to the same degree. Thus, we interpret this lack of difference in SOC-derived MBC and SOC-derived CO_2_ between glucose- and pine-amended soils as evidence that priming is likely not impacting our results.

As expected, burning does not affect either microbial biomass or respiration in underlying mineral soil, which, owing to its greater depth below the soil surface, receives less heat during a fire than surface horizons. For example, maximum soil temperatures following the 120 s heat dose treatments reached an average of 337 °C at 5 cm below the soil surface in Histosol cores and 163 °C at O horizon-mineral soil interface in Gleysols. At the base of the cores, average maximum soil temperatures were 57 °C and 36 °C in Histosol and Gleysol cores (Johnson et al. 2024). This difference in temperature with depth suggests that fire has a substantially reduced effect on C cycling in subsurface horizons and that *in situ* measurements of decreased post-fire soil respiration are likely primarily driven by changes in surficial soil horizons, at least in the short-to intermediate term after the fire.

In the few studies that have measured soil CO_2_ efflux in the hours to days following fire, a sharp increase in soil respiration is often observed, which may be caused by the mineralization of fire-generated microbial and plant necromass (Bárcenas-Moreno et al. 2011, Masyagina et al. 2016, Köster et al. 2024). By three weeks post-burn treatment in these samples, this pulse of respiration had subsided and the relative rate of mineralization (per g remaining C) in burned soils dropped below rates in unburned soils (Johnson et al. 2024), suggesting that most of any newly available C in necromass resulting from the fire had largely been degraded by the surviving microbes by the time this study was initiated. For this study, soil subsamples were homogenized 24 days post-fire before beginning the 4-day CUE incubation. After homogenizing the soil, respiration rates in the burned soils were still lower than rates in unburned soils, which suggests that spatial accessibility alone was not constraining soil respiration rates. Instead, lower mineralization rates of remaining C in burned soils compared to unburned soils are likely primarily driven by fire-induced shifts in C chemistry, as previously suggested (Johnson et al. 2024).

### 4.2. The decrease in glucose-specific CUE with burning correlates with an increase in predicted mean 16S rRNA gene copy number

CUE_glucose_ was lower in burned *vs*. unburned organic horizon soils from both soil types, which was consistent with our first hypothesis that burning would reduce CUE. Decreased CUE can be driven by increased respiration and/or decreased microbial growth. In this study, the biggest driver is decreased microbial growth – burning had small to no effects on glucose-derived respiration (Figure 4B), which is evidence that the microbial community in burned soils is still active and degrading the added glucose. At the same time, we observed substantially lower incorporation of C from glucose into microbial biomass (*i*.*e*., microbial growth) in burned soils, hence the decrease in CUE_glucose_ in burned organic horizon soils.

Previous work has demonstrated a general increase in the relative abundance of fast growing bacterial taxa following fire, which was positively correlated with burn severity (Johnson et al. 2023). The heat-induced production of necromass during a fire may stimulate an increase in fast-growing taxa dependent upon simple C substrates that are easily accessible to microbes (*i.e.*, not sorbed to mineral surfaces). Our observed decrease in CUE_glucose_ in burned soils (Figure 5B) and decreasing CUE_glucose_ with increasing maximum soil temperature (Figure 6A) are consistent with a bloom of fast-growing, inefficient taxa post-fire. This was supported by our observations of an exponential decrease in CUE_glucose_ with increasing predicted weighted mean 16S rRNA gene copy number (Figure 6B), which has been shown to correlate with microbial maximum potential growth rates (Roller et al. 2016, Johnson et al. 2023). Thus, while putatively fast-growing taxa in burned soils seem to be mineralizing glucose in burned soils at a rate similar to or faster than unburned microbial communities, these fast growers appear to be less efficient at incorporating C from glucose into biomass.

Our finding of decreased CUE_glucose_ but no consistent change in CUE_pine_ in burned soils is also supportive of fast-growing taxa preferentially utilizing simpler substrates like glucose over more chemically complex substrates such as pine. There were small increases in glucose-derived respiration with burning (Figure 4), which is consistent with glucose degradation by a bloom of fast-growing, less efficient taxa post-fire.

It is possible that burn-induced changes in the soil environment are also shaping post-fire CUE. Elevated soil pH following burning may have contributed to decreased CUE_glucose_ (Figure S1), although the trend does not clearly support any of the previously noted observed relationships between pH and CUE (Sinsabaugh et al. 2016, Silva-Sánchez et al. 2019, Jones et al. 2019, Schroeder et al. 2024). Furthermore, despite a wide range of measured pH values (from 4 to 7), the lack of a clear effect of pH on CUE_glucose_ in Gleysols (Figure S1) suggests that pH may be a relatively minor or an indirect and inconsistent driver of post-fire CUE.

The effects of burning on CUE_glucose_ in underlying mineral soils are more subtle than those in organic horizons (Figure 5), which is consistent with previous work finding more muted effects of burning on soil properties and microbial communities with depth (Hartford and Frandsen 1992, O’Donnell et al. 2011, Knicker et al. 2013, Pressler et al. 2019, Johnson et al. 2024). This can be attributed to the lower temperatures reached during fire in underlying mineral soils (Johnson et al. 2024) due to the insulative properties of soil and a resulting smaller impact of fire on available C substrates and microbial community composition. This close relationship between heat exposure and CUE is reflected in the significant exponential decrease in CUE_glucose_ with increasing maximum soil temperatures reached during the burns in both the organic horizon and mineral soils (Figure 6A).

The net effect of an increased mean community growth rate on CUE will depend upon the persistence of the fast-growing fraction of the microbial community. Although the impact of burning on CUE_glucose_ in mineral soils was small, previous work has shown that the enrichment of fast-growing taxa can persist for at least one year following fire in both organic and mineral soils from this region (Johnson et al. 2023). If the observed decreases in CUE_glucose_ in organic and mineral soil following burning is driven primarily by a shift in microbial community composition to more fast-growing, less-efficient taxa, then it is possible that the return of CUE to pre-burn levels could require a year or more. Furthermore, the negative relationship between CUE_glucose_ and maximum soil temperatures suggests that hotter fires may have larger negative impacts on CUE, and thus, larger impacts on soil C cycling in the months to years following fire. These changes in CUE could have important implications for C stocks. If microbes process C differently after a burn by incorporating proportionally less C into biomass, that would also result in less necromass when the microbes die. Because of the close interactions between living microbes and mineral surfaces, their necromass may disproportionately contribute to mineral-associated organic matter, which is thought to be more persistent (Cotrufo et al. 2013). Thus, the timescale over which this effect on CUE persists will be important in determining the scale of any potential downstream effects on soil C stocks. Given that our past work suggests that enrichment of fast-growers is limited to less than five years post-fire in this region (Johnson et al. 2023), the long-term effects of this change on soil C stocks may be limited.

### 4.3. Pine-specific CUE is lower than glucose-specific CUE in unburned soils but the effect of burning on pine-specific CUE is small

CUE_pine_ was lower in unburned organic horizon soils than CUE_glucose_ (Figure 5), supporting our second hypothesis that CUE of the more complex C substrate would be lower. However, unlike with CUE_glucose_, we saw much smaller to no shifts in CUE_pine_ with burning, which was contrary to our first and central hypothesis that burning would decrease CUE. There are three possible explanations for these findings. First, it is possible that burning truly has little to no effect on the CUE of more complex substrates like pine. Our results suggest that the portion of the microbial community responsible for degradation of more complex substrates, such as pine, is less affected by burning than fast growing taxa using simpler substrates, such as glucose. This is supported by the lack of effect of burning on pine-derived respiration (Figure 4A) or the incorporation of C from pine into microbial biomass (Figure 3A).

An alternative explanation for the lack of effect of burning on CUE_pine_ is that the 4-day incubation was not long enough to accurately estimate CUE_pine_, due to the higher complexity of pine relative to glucose. If the incubation length was too short to allow for all the enzymatic steps required to break down a large portion of the added pine into components that can be taken up by a cell, then the treatment effects of burning on CUE_pine_ may not be evident. Less than 10% of added pine C was decomposed during this incubation on average. Thus, the ^13^C signal from this pine may have been too small to result in measurable differences between burned and unburned soils. Results for seven of 54 of the pine-amended soils showed negative values for substrate-derived MBC (reflecting statistical error around 0), compared to one of 54 soils amended with glucose. While burning did not have a significant effect on pine-derived microbial biomass, there were small, albeit non-significant, decreases in pine-derived microbial biomass in the soils exposed to the 120 s heat dose treatment compared to the unburned soil (Figure 3A). Choosing an appropriate incubation length for estimating CUE is notoriously difficult, because while all CO_2_ emissions during the incubation can be captured, microbial biomass turnover over the course of the incubation risks underestimating true net incorporation of substrate C into biomass. Furthermore, because we used true field replicates, with each core representing a completely separate field site, variation is high. Additionally, because we used realistic burns in intact cores (rather than heating each soil to a controlled temperature, for example), there was substantial variability in how each burn proceeded within each burn treatment. Hence, it is possible that with either a longer incubation time period, a stronger ^13^C label of labelled pine, or less variability across soil cores and burns, these differences with burning would become more pronounced/detectable.

Another possible explanation for the lack of significant effect of burning on CUE_pine_ is the potential impact of homogenizing soil samples on microbial community composition (West and Whitman 2022). Soils were homogenized before being amended with pine or glucose, and we expect that while homogenization would impact both bacteria and fungi, it might have had a larger impact on fungal communities than bacterial communities, due to the physical disruption of hyphal networks. If fungal abundance and CUE are indeed positively correlated, then disrupting fungal hyphae via homogenization may have caused a decrease in CUE across all treatments that overshadowed the effect of burning on CUE_pine_. Because many soil fungi decompose woody biomass, it is possible that the disruption of soil fungi as part of the experimental process may particularly affect our ability to accurately measure CUE_pine_ in burned or unburned soils. However, more work is needed to understand the relationship between fungal abundance and CUE in boreal forest soils before we will be able to untangle the impact of fire on this relationship.

Overall, the baseline differences between CUE_pine_ and CUE_glucose_ were smaller than burn-induced shifts in CUE (where significant), supporting our third hypothesis that substrate-specific differences would be smaller than changes in CUE driven by burning. This suggests that burn-induced changes in microbial community composition and function do play an important role in mediating post-fire C cycling.

### 4.4. No evidence of homefield advantage for microbial communities from pine forests

Our final hypothesis, that the difference between CUE_glucose_ and CUE_pine_ would be smaller in Gleysols than in Histosols, was not supported by our observations. We hypothesized that microbial communities from the Gleysol sampling sites may have a “home field advantage” (Fanin et al. 2016) in degrading pine because these sampling sites were largely dominated by *Pinus banksiana* – the same genus as *Pinus strobus*, which was the source of the pine amendments. Instead, the difference between CUE_glucose_ and CUE_pine_ was even larger in the organic horizons of Gleysols than in Histosols for unburned soils. This suggests that the microbial communities from the Gleysols are not meaningfully more efficient at pine degradation than the communities from the Histosols. Alternately, there could be a large enough structural or chemical difference in *Pinus strobus* and *Pinus banksiana* plant material to negate a home field advantage effect.

## Summary and Future Directions

Burning generally decreases soil microbial community CUE by causing a decrease in microbial growth, with increases in respiration playing a smaller role. Burning had little impact on the rate of respiration of C derived from either of the added substrates, suggesting that decreases in post-fire soil respiration are more likely driven by changes in C chemistry and the concomitant change in microbial efficiency, which is then reflected in which microbes are active, than by changes in total microbial biomass. Instead, despite potentially large reductions in total size, the microbial community in burned soil continues to decompose added C substrates, albeit at a less efficient rate. Larger decreases in the CUE of glucose than of pine, a more complex substrate, also support the possibility that the post-fire chemical composition of remaining SOC may be a more important driver of post-fire soil C fluxes than microbial biomass or community composition.

These results highlight the influence of environmental factors on soil CUE and support previous calls for a more nuanced representation of CUE in soil C models. The observed relatively strong negative correlation (R^2^ = 0.51) between CUE and weighted mean predicted 16S rRNA gene copy number was intriguing to us because of its potentially important implications for modelling. If weighted mean predicted 16S rRNA gene copy number is a viable proxy for estimating CUE, it could dramatically reduce the time and resources required to experimentally measure CUE at multiple timepoints post-fire. Many soil C cycling models rely on CUE as a central parameter (Allison 2025), and recent work indicates that CUE is critically important in driving global models of SOC stocks (Tao et al. 2023). However, measuring CUE is laborious, technical, and expensive, and laboratory methods used to measure CUE can be quite variable (Geyer et al. 2019, He et al. 2024). Thus, global-scale datasets of empirically-derived CUE are fundamentally limited. In contrast, 16S rRNA gene amplicon sequencing data are increasingly common, standardized, and publicly available. We are excited about the potential for leveraging globally-extensive sequencing datasets to estimate CUE in biogeochemical models, or at least to scale differences in CUE across different samples, environments, and treatments across different geographic scales. Given the relationship between CUE, weighted mean predicted 16S rRNA gene copy number, and fast-growing taxa, sequencing data could also potentially inform the partitioning of fast / low-CUE and slow / high-CUE soil microbial pools in microbially-explicit models.

However, this study is temporally limited, representing a single timepoint of post-burn CUE, and there are numerous ways in which fire may affect CUE – *e.g.* changes in soil temperature, shifts in plant inputs – that are not explored here and may be important for understanding post-fire CUE. Future work will help elucidate how substrate-specific CUE may vary in the months to years post-fire and which mechanisms contribute to the recovery of CUE to pre-burn levels.

## Acknowledgements

We thank D. Letourneau and M.A. Parisien at Natural Resources Canada for assistance with field work; J. Morin, T.J. Little, and other Wood Buffalo National Park staff for support in conducting this research (Permit WB-2022-41998); E. Lazarcik and K. Bourne at the USDA FS Forest Products Laboratory for facilitating the burn simulations; Z. Freedman for assistance with headspace gas CO_2_-C analysis; T. Berry for help with ^13^C isotope measurements and calibrations; K. Kruger and A. Hutka for assistance with laboratory work; N. Zeba, M. Sikora, and J. Woolet for producing the ^13^C-labelled pine material; and H. Reid for assistance with data acquisition. This study was funded by a U.S. Department of Energy grant to TW (DE-SC0021022).

## Author contribution statement

**Dana Johnson**: Conceptualization, Methodology, Formal analysis, Investigation, Writing – original draft, Visualization. **Kara Yedinak**: Methodology, Resources, Writing – review and editing. **Thea Whitman**: Funding acquisition, Conceptualization, Methodology, Resources, Writing – review and editing.

