## Supplemental Information for "Burn-induced decreases in soil microbial carbon use efficiency vary across soil types and substrates"

Table of Contents	Page

**Supplementary Tables**

Table S1. Site locations and characteristics	2

Table S2. Horizon thickness, C and N concentrations, and pH	2

Table S3. Mass of amendment C added to soil	3

**Supplementary Figures**

Figure S1. Microbial CUE_glucose_ vs. soil pH	4

Figure S2. Microbial CUE_pine_ vs. weighted mean gene copy number	4

**Supplementary Methods**

Eq. S1. Adjusted DOC concentrations of fumigated solutions	5

Eq. S2. Adjusted DOC concentrations of non-fumigated solutions	5

Eq. S3. Total microbial biomass C	5

Eq. S4. Atom % of ^13^C in MBC with labeled amendments	5

Eq. S5. Atom % of ^13^C in MBC with unlabeled amendments	5

Eq. S6. Substrate-derived MBC as fraction of total MBC	6

Eq. S7. Mass of substrate-derived MBC	7

Eq. S8. Substrate-derived CO_2_ as fraction of total CO_2_	7

Eq. S9. Mass of substrate-derived CO_2_	8

Eq. S10. CUE	8

Eq. S11. Metabolic quotient, qCO_2_	8

**Supplementary Tables**

| Table S1. Site locations and characteristics. | | | | | |
| --- | --- | --- | --- | --- | --- |
| **Site ID** | **Latitude** | **Longitude** | **Soil texture class** | **Soil type** | **Dominant tree species** |
| 1 | -112.26 | 59.49 | Loamy Sand | Gleysol | *Pinus banksiana* |
| 2 | -112.41 | 59.40 | Sand | Gleysol | *Pinus banksiana* |
| 3 | -112.49 | 59.33 | Sand | Gleysol | *Pinus banksiana* |
| 5 | -112.39 | 59.41 | Sandy Loam | Gleysol | *Pinus banksiana* |
| 8 | -112.48 | 59.36 | Silt Loam | Gleysol | *Pinus banksiana* |
| 9 | -112.41 | 59.38 | Sand | Gleysol | *Pinus banksiana* |
| 4 | -112.49 | 59.31 | Organic | Histosol | *Picea* spp. |
| 6 | -112.25 | 59.51 | Organic | Histosol | *Picea* spp. |
| 7 | -112.36 | 59.44 | Organic | Histosol | *Picea* spp. |
| 10 | -112.42 | 59.39 | Organic | Histosol | *Picea* spp. |
| 11 | -112.38 | 59.43 | Organic | Histosol | *Picea* spp. |
| 12 | -112.49 | 59.33 | Organic | Histosol | *Picea* spp. |

| Table S2. Soil horizon thickness, C and N concentrations, and pH (mean ± standard deviation) in burned and unburned soils. | | | | | |
| --- | --- | --- | --- | --- | --- |
| **Soil type** | **Burn duration treatment (s)** | **Soil horizon thickness (cm)** | **C conc. (%)** | **N conc. (%)** | **pH** |
| Histosol | 0 | 10 ± 0 | 39.7 ± 7.4 | 1.3 ± 0.2 | 6.6 ± 0.74 |
|  | 30 | 9.5 ± 0.71 | 39.2 ± 5.2 | 1.4 ± 0.2 | 7.3 ± 0.60 |
|  | 120 | 8.7 ± 0.88 | 37.6 ± 7.3 | 1.5 ± 0.3 | 7.7 ± 0.65 |
| Gleysol, O horizons | 0 | 2.4 ± 0.78 | 18.9 ± 8.3 | 0.6 ± 0.3 | 4.6 ± 0.19 |
|  | 30 | 2.4 ± 0.69 | 21.1 ± 9.3 | 0.7 ± 0.3 | 5.0 ± 0.41 |
|  | 120 | 2.7 ± 0.96 | 26.6 ± 12.1 | 1.0 ± 0.5 | 6.1 ± 1.1 |
| Gleysol, mineral horizons | 0 | 7.5 ± 0.78 | 1.5 ± 0.5 | 0.06 ± 0.02 | 4.8 ± 0.45 |
|  | 30 | 7.4 ± 0.71 | 1.5 ± 0.5 | 0.06 ± 0.02 | 4.9 ± 0.48 |
|  | 120 | 7.2 ± 0.93 | 1.6 ± 0.5 | 0.06 ± 0.02 | 5.0 ± 0.48 |

| Table S3. Mass of amendment C added to soil for incubation and mass of chloroform used during microbial biomass C measurement | | | |
| --- | --- | --- | --- |
| **Addition to soil** | **Soil type** | **C added**  **(mg per g dry soil)** | **C added**  **(percent of total C)** |
| Glucose | Histosol | 0.4 ± 0.002 | 0.11% |
|  | O horizon, Gleysol | 0.4 ± 0.005 | 0.23% |
|  | Mineral soil, Gleysol | 0.4 ± 0.0008 | 2.9% |
| Pine | Histosol | 1.0 ± 0.005 | 0.27% |
|  | O horizon, Gleysol | 1.0 ± 0.01 | 0.57% |
|  | Mineral soil, Gleysol | 1.0 ± 0.002 | 7.2% |
| Chloroform* | Histosol | 109 ± 2 | 29% |
|  | O horizon, Gleysol | 120 ± 55 | 75% |
|  | Mineral soil, Gleysol | 17 ± 0.4 | 120% |

* Traces of chloroform were removed from the extracted solution by bubbling with lab air before analysis

**Supplementary Figures**

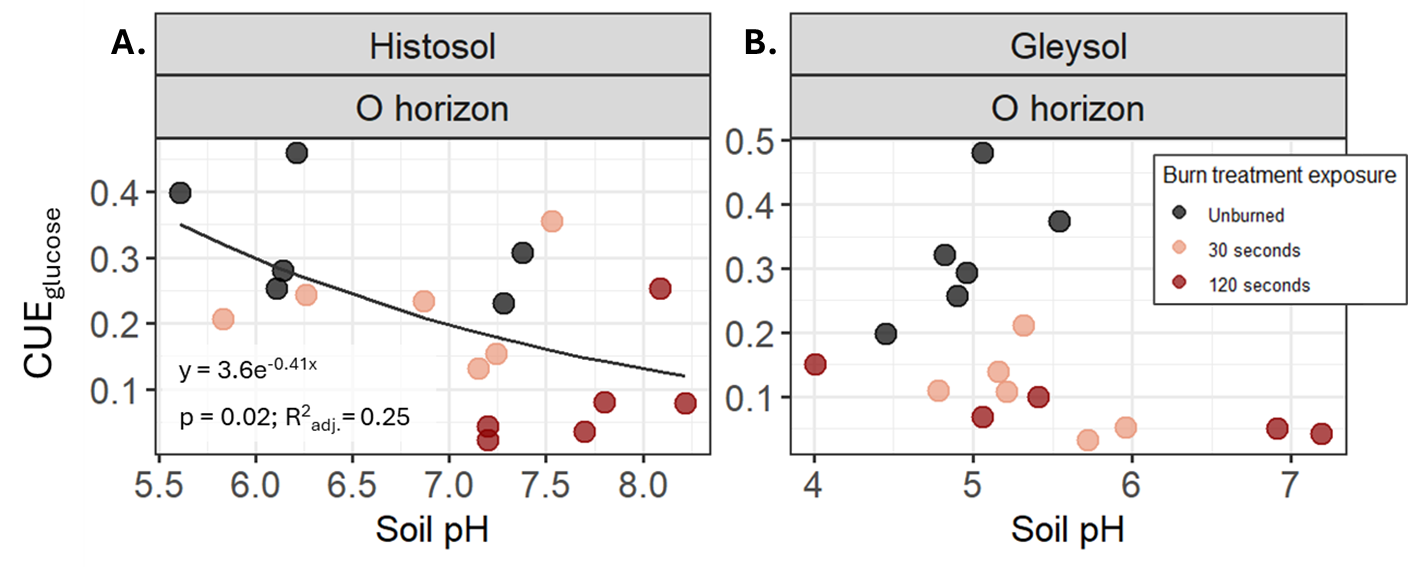

*Figure S1. Soil pH vs. CUE_glucose_ in Histosol (A) and Gleysol (B) O horizon soils. We used a non-linear regression to test for a significant correlation between maximum temperature and CUE.*

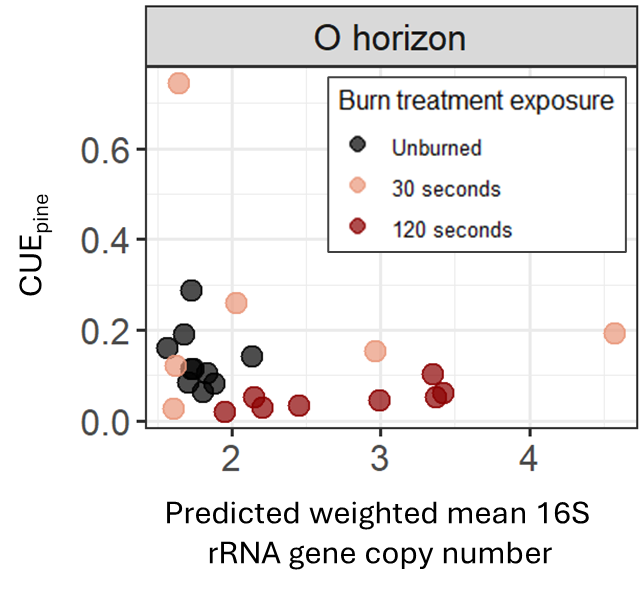

*Figure S2. Predicted weighted mean 16S rRNA gene copy number vs. CUE_pine_ in organic horizon soils. We used a non-linear regression to test for a significant correlation between weighted mean gene copy number and CUE. Soil type did not have a significant effect, so results for Histosols and Gleysols are plotted together.*

**Supplementary Methods**

Carbon use efficiency calculations and derivatizations

Adjust dissolved organic C (DOC) concentrations following Equations S1 & S2:

DOC_F,adj._ = DOC_F_ – DOC_F,blank_	Eq. S1

DOC_NF,adj._ = DOC_NF_ – DOC_NF,blank_	Eq. S2

where DOC_F_ and DOC_NF_ represent the total dissolved organic carbon (µg C g^-1^ dry soil) from fumigated (F) and non-fumigated (NF) soils, respectively, and DOC_F,adj._ and DOC_NF,adj._ represent blank-adjusted total dissolved organic carbon.

Calculate total microbial biomass C using Equation S3:

$\text{MBC}= \text{DOC}_{\text{F, adj}}- \text{DOC}_{\text{NF,adj}}$	Eq. S3

where MBC represents total microbial biomass C (µg C g^-1^ dry soil).

Calculating the atom % of ^13^C in MBC from soils with labelled and unlabeled amendments using Equations S4 and S5, respectively:

$\text{at\% }\text{MBC}_{\text{soil+LA}}= \frac{\left( \text{at\%}\text{DOC}_{\text{F,LA}} \times\left[ \text{DOC}_{\text{F,LA}} \right] \right)-(\text{at\%}\text{DOC}_{\text{NF,LA }}\times\left[ \text{DOC}_{\text{NF,LA}} \right])}{(\text{DOC}_{\text{F,LA}}-\text{DOC}_{\text{NF,LA}})}$	Eq. S4

$\text{at\% }\text{MBC}_{\text{soil+UA}}= \frac{\left( \text{at\%}\text{DOC}_{\text{F,UA}} \times\left[ \text{DOC}_{\text{F,UA}} \right] \right)-(a\text{t\%}\text{DOC}_{\text{NF,UA }}\times\left[ \text{DOC}_{\text{NF,UA}} \right])}{(\text{DOC}_{\text{F,UA}}-\text{DOC}_{\text{NF,UA}})}$	Eq. S5

where at% MBC_soil+LA_ represent the atom % of ^13^C in total MBC from soils with the labelled amendment (LA), and at% DOC_F,LA_ and at% DOC_NF,LA_ represent the atom % of ^13^C in DOC from soils with the labelled amendment that were fumigated or non-fumigated, respectively. DOC_F,LA_ and DOC_NF,LA_ represent the DOC (µg C g^-1^ dry soil) from fumigated and non-fumigated soils that were first amended with the labelled substrate. At% MBCsoil+_UA_ represents the atom % of ^13^C in total MBC from soils with the unlabeled amendment (UA), and at% DOC_F,UA_ and at% DOC_NF,UA_ represent the atom % of ^13^C in DOC from soils with the unlabeled amendment that were fumigated or non-fumigated, respectively. DOC_F,UA_ and DOC_NF,UA_ represent the DOC (µg C g^-1^ dry soil) from fumigated and non-fumigated soils that were first amended with the unlabeled substrate.

Calculate the fraction of total MBC that is substrate-derived using Equation S6:

${}{f_{MBC,LA}= \frac{(\text{at\%}\text{MBC}_{\text{soil+LA}}-\text{at\%}\text{MBC}_{\text{soil+UA}})}{(\text{at\%LA}-\text{at\%UA})}}$	Eq. S6

where f_MBC,LA_ is the fraction of total MBC derived from the labelled amendment, and at% LA and at% UA is the atom % of ^13^C in the labelled and unlabeled amendments, respectively.

Derivation of Equation S6

Starting equations:

$${at\%MBC}_{soil+LA}=f_{MBC,LA}\times at\%LA+ f_{MBC,soil}\times at\%soil$$

$${at\%MBC}_{soil+UA}=f_{MBC,UA}\times at\%UA+ f_{MBC,soil}\times at\%soil$$

Assumption: $f_{MBC,LA}\approx f_{MBC,UA}$

Derivation steps

1. $f_{MBC,soil}\times at\%soil={at\%MBC}_{soil+LA}-f_{MBC,LA}\times at\%LA$
2. $f_{MBC,soil}\times at\%soil={at\%MBC}_{soil+UA}-f_{MBC,UA}\times at\%UA$
3. $f_{MBC,soil}\times at\%soil={at\%MBC}_{soil+UA}-f_{MBC,LA}\times at\%UA$
4. ${at\%MBC}_{soil+LA}-f_{MBC,LA}\times at\%LA={at\%MBC}_{soil+UA}-f_{MBC,LA}\times at\%UA$
5. ${at\%MBC}_{soil+LA}-{at\%MBC}_{soil+UA}=f_{MBC,LA}\times at\%LA-f_{MBC,LA}\times at\%UA$
6. ${at\%MBC}_{soil+LA}-{at\%MBC}_{soil+UA}=f_{MBC,LA}\times(at\%LA-at\%UA)$
7. ${}{f_{MBC,LA}= \frac{(\text{at\%}\text{MBC}_{\text{soil+LA}}-\text{at\%}\text{MBC}_{\text{soil+UA}})}{(\text{at\%LA}-\text{at\%UA})}}$

Calculating the total substrate-derived MBC (µg C g^-1^ dry soil) using Equation S7:

$\mathrm{MBC}_{\mathrm{LA}}=MBC\times f_{MBC,LA}$	Eq. S7

where MBC_LA_ represents substrate-derived MBC (µg C g^-1^ dry soil).

Calculating the fraction of total CO_2_ that is derived from the labelled substrate using Equation S8:

$f_{CO2, LA}=\frac{\delta^{13}C_{CO2,soil+LA}- \delta^{13}C_{CO2,soil+UA}}{\delta^{13}C_{\mathrm{LA}}-\delta^{13}C_{\mathrm{UA}}}$	Eq. S8

where f_CO2,LA_ is the fraction of total CO_2_ derived from labelled amendment, δ^13^C_CO2,soil+LA_ and δ^13^C_CO2,soil+UA_ represent the δ^13^C values of CO_2_ from soils and the labelled or unlabeled amendment, respectively, and δ^13^C_LA_ and δ^13^C_UA_ represent the δ^13^C values of the labelled and unlabeled amendments, respectively.

Derivation of Equation S8

Starting equations:

$$\delta^{13}C_{CO2, soil+LA}=f_{CO2,soil}\times\delta^{13}C_{soil}+f_{CO2,LA}\times\delta^{13}C_{LA}$$

$$\delta^{13}C_{CO2, soil+UA}=f_{CO2,soil}\times\delta^{13}C_{soil}+f_{CO2,UA}\times\delta^{13}C_{UA}$$

Assumption: $f_{LA}=f_{UA}$

Where δ^13^C_CO2,soil+LA_ and δ^13^C_CO2,soil+UA_ represent the δ^13^C of CO_2_ from soils with the labelled or unlabeled amendment, respectively; f_CO2,soil_ and δ^13^C_soil_ represent the fraction of total CO_2_-C derived from soil and the δ^13^C signature of the soil, respectively; f_CO2,LA_ and f_CO2,UA_ represent the fraction of total CO_2_-C derived from the labelled and unlabeled amendments, respectively; and δ^13^C_LA_ and δ^13^C_UA_ represent the δ^13^C signature of the labelled and unlabeled amendment, respectively.

Calculate total substrate-derived CO_2_ (µg C g^-1^ dry soil) using Equation S9:

$\mathrm{CO}_{2 LA}=\mathrm{CO}_{2 total}\times f_{CO2,LA}$	Eq. S9

where CO_2 LA_ represents substrate-derived CO_2_ (µg C g^-1^ dry soil), and CO_2 total_ represents total CO_2_ respired (µg C g^-1^ dry soil).

Calculate CUE using Equation S10:

$\text{CUE}= \frac{\mathrm{MBC}_{\mathrm{LA}}}{(\mathrm{MBC}_{\mathrm{LA}}+\mathrm{CO}_{2,LA})}= \frac{substrate derived MBC}{(substrate derived MBC+substrate derived \mathrm{CO}_{2})}$	Eq. S10

Calculate the metabolic quotient, qCO_2_, using Equation S11:

$\mathrm{qCO}_{2}=\frac{\mathrm{CO}_{2,\mathrm{LA}}}{\mathrm{MBC}_{\mathrm{LA}}}$	Eq. S11
